# Validating Paired-end Read Alignments in Sequence Graphs

**DOI:** 10.1101/682799

**Authors:** Chirag Jain, Haowen Zhang, Alexander Dilthey, Srinivas Aluru

## Abstract

Graph based non-linear reference structures such as variation graphs and colored de Bruijn graphs enable incorporation of full genomic diversity within a population. However, transitioning from a simple string-based reference to graphs requires addressing many computational challenges, one of which concerns accurately mapping sequencing read sets to graphs. Paired-end Illumina sequencing is a commonly used sequencing platform in genomics, where the paired-end distance constraints allow disambiguation of repeats. Many recent works have explored provably good index-based and alignment-based strategies for mapping individual reads to graphs. However, validating distance constraints efficiently over graphs is not trivial, and existing sequence to graph mappers rely on heuristics. We introduce a mathematical formulation of the problem, and provide a new algorithm to solve it exactly. We take advantage of the high sparsity of reference graphs, and use sparse matrix-matrix multiplications (SpGEMM) to build an index which can be queried efficiently by a mapping algorithm for validating the distance constraints. Effectiveness of the algorithm is demonstrated using real reference graphs, including a human MHC variation graph, and a pan-genome de-Bruijn graph built using genomes of 20 B. *anthracis* strains. While the one-time indexing time can vary from a few minutes to a few hours using our algorithm, answering a million distance queries takes less than a second.

**2012 ACM Subject Classification:** Mathematics of computing → Paths and connectivity problems; Applied computing → Computational genomics

## 1 Introduction

Owing to continuous technological and algorithmic advancements in genomics during the past four decades, whole-genome sequencing has now become ubiquitous, leading to an explosive growth in genome databases. Despite this progress, the current human genome reference (GRCh38) is primarily derived from a single individual [14, 38]. Many recent studies have demonstrated improved variant analysis using a graph-based reference while accounting for population diversity. Accounting for this diversity is especially critical in polymorphic regions of genomes [10, 17]. Realizing this paradigm shift from a simple linear reference to a graph-based reference, however, requires addressing several open computational challenges [5], one of which concerns designing robust algorithms for mapping reads to graph-based references. Accuracy of read mapping is critical for downstream biological analyses.

Designing provably good algorithms for approximate sequence matching to graphs, using both index-based and alignment-based approaches, remains an active research area. A few recent works have investigated extending Burrows-Wheeler-Transform-based indexing to sequence DAGs [41] and de-Bruijn graphs [2, 29, 40]. Similarly, there exist studies that have explored extension of the classic sequence-to-sequence alignment routines to graphs [19, 20, 30, 36]. In our recent work [19], we presented new complexity results and algorithms for the alignment problem using general sequence-labeled graphs. The results show that a sequence (of length *m*) can be aligned to a labeled directed graph *G*(*V, E*) in *O*(|*V*| + *m* |*E*|) time, using commonly used scoring functions, while allowing edits in the query but not graph labels. However, a general string to graph pattern matching formulation is only good for mapping single-end reads or single-molecule sequencing reads, and does not account for pairing information.

Paired-end sequencing provides information about the relative orientation and genomic distance between the two reads in a pair. When reads originate from repetitive regions, this information is valuable for pruning large number of false candidates [4]. Popular short read mapping tools for linear references, e.g., BWA-mem [24] and Bowtie2 [22], therefore, enforce these constraints in a read pair to guide the selection of the true mapping locus. Using a linear reference, calculating gap between two mapping locations is just a simple subtraction operation. However, it still remains unclear how to efficiently validate the constraints using large non-linear graph-based references and read sets.

Several sequence to graph aligners have been developed in recent years to map reads to variation graphs [11, 12, 18, 21, 28, 35, 37], de-Bruijn graphs [16, 25, 26] and splicing graphs [1, 8]. Readers are referred to review articles, e.g., [5, 34] for an expanded list of the tools. Among these tools, Graph-Aligner [35], vg [12], deBGA [26], HISAT2 [21] and HLA-PRG [11] support paired-end read mapping. However, all of these use heuristics to measure the observed insert size between the two reads in a pair, mainly due to lack of associated provably-good graph-based algorithms. A popular heuristic adopted by the tools is to do the computation while assuming a linear ordering of vertices (e.g., topological order). However, it can produce misleading results in complex variation-rich graph regions.

In this work, we provide the first mathematical formulation of the problem of validating paired-end distance constraints in sequence graphs, and propose an exact algorithm to solve it that is also practical. The proposed algorithm exploits sparsity in sequence graphs to build an index, which can be queried quickly using a simple lookup during the read mapping process. On the performance side, we provide formal arguments to shed light on why our indexing procedure is efficient with regards to the computation time and storage requirements. We show the practical significance of our algorithm using LRC_KIR and MHC variation graphs derived from the human genome, as well as pan-genomic de Bruijn graphs of *Bacillus anthracis* strains.

## 2 Problem Formulation

### Definition 1

*Sequence Graph: A sequence graph G*(*V, E*) *is a directed graph with vertices V and edges E, where each vertex v* ∈ *V is labeled with a character from alphabet* Σ.

Sequence graph is typically defined as a directed graph with either string or character labeled vertices because converting one form into the other is straightforward. In addition, commonly used graph formats, such as de-Bruijn graphs, bi-directional de-Bruijn graphs, overlap graphs or variation graphs can be converted into sequence graphs with at most a constant factor increase in vertex or edge set sizes. While single-end read alignments can be judged by their alignment scores alone, a valid paired-end read alignment over a sequence graph should satisfy the expected paired-end distance constraints and orientation. As insert size can vary within a range, let *d*_1_ and *d*_2_ denote the minimum and maximum allowed values of the inner distance between the reads within a pair (see Figure 1).

**Figure 1.**
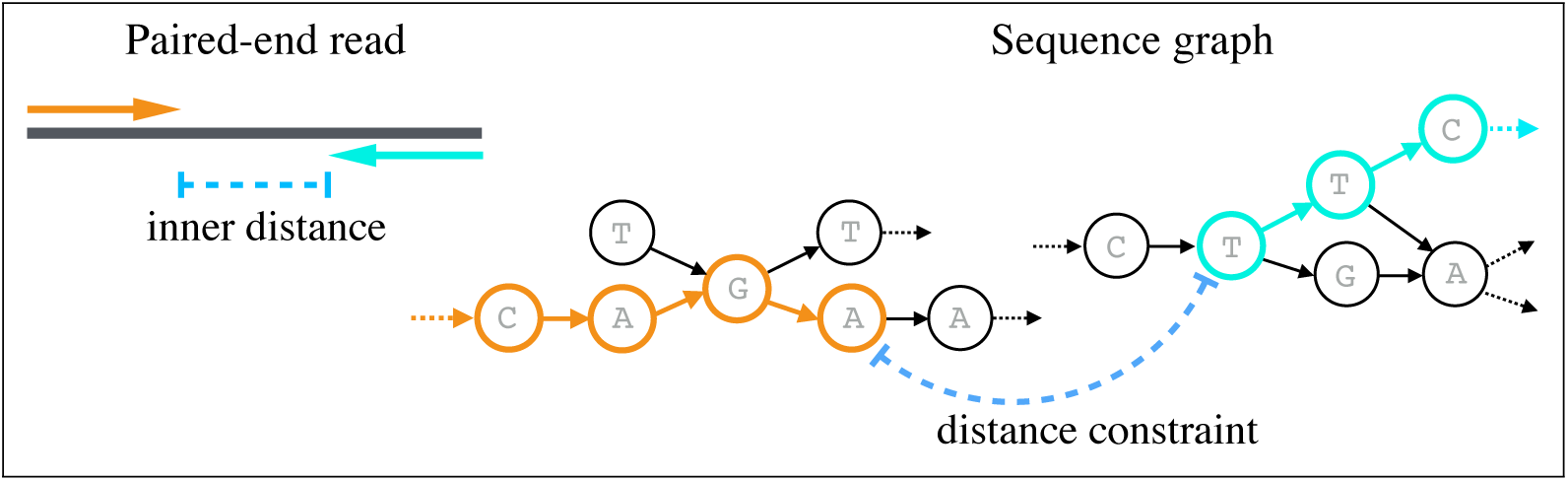
Visualizing distance constraints while mapping paired-end reads to sequence graphs.

### Definition 2

*Paired-end Validation Problem: Suppose two reads r*_1_, *r*_2_ *in a pair are independently mapped to a sequence graph G*(*V, E*) *using their positive and negative strands respectively. Let v*_1_ *be the vertex where a path to which r*_1_ *(+ve strand) is mapped ends, and v*_2_ *be the vertex where a path to which r*_2_ *(-ve strand) is mapped starts; then we refer to this pair of paths as a valid paired-end read mapping if and only if there exists a path from v*_1_ *to v*_2_ *of length d* ∈ [*d*_1_, *d*_2_].

The above problem definition is based on the assumption that a fragment, from which a read pair is sequenced, can align to any (cyclic or acyclic) path in the input sequence graph. In this work, we focus on designing an efficient algorithm that can quickly answer the above path queries for any two given vertices in the graph. Typically, there are multiple mapping candidates to evaluate for each read pair, especially if a read is sequenced from repetitive regions of a genome. In addition, a typical read set in a genomic study may contain millions or billions of reads. Therefore, validating the distance queries quickly using an appropriate indexing scheme is desirable.

## 3 Related Problems in Graph Theory

Computing all-pairs shortest paths in *G*(*V, E*) may help to identify true-positives or true-negatives, but only for those vertex pairs whose shortest distance ≥ *d*_1_. No conclusion can be drawn when the shortest distance between two vertices is *< d*_1_, as a valid path need not be the shortest path. In addition, computing all-pairs shortest paths is expensive, and may not provide the desired scalability [7]. If *d*_1_ = *d*_2_, the formulated problem becomes a special case of the exact-path length problem [33], with all edge weights set to 1. The exact-path length problem determines if a path of a specified distance exists between two vertices in a weighted graph. An extension of this problem, referred to as the gap-filling problem [39], has been explored in the context of genome assembly using paired-end or mate pair read sets. Although the exact-path length problem has been shown to be𝒩𝒫-complete [33], we will demonstrate a simple and practical polynomial-time algorithm for our problem with unweighted edges. Finally, if *d*_1_ = 0 and *d*_2_ = |*V*|, then our problem is equivalent to determining transitive closure of a graph [32]. In our case, however, we expect *d*_2_ ≪|*V*|.

Our approach is based on an indexing strategy where we pre-compute a boolean index matrix, which has a 1 for each vertex pair that satisfies the distance constraints (Section 4). Computing the index requires polynomial operations, and paired-end distance queries can be computed quickly using index lookups during the read mapping process. Before describing the algorithm, we first discuss a trivial pseudo-polynomial time algorithm to solve the paired-end distance validation problem. It is based on a well-known algorithm used to solve the intractable subset-sum problem.

### A Pseudo-polynomial Time Algorithm

The problem of validating distance constraints between two vertices can be solved using dynamic programming. Assume *s* ∈*V* is the source vertex from where we need to query paths of length *d* ∈ [*d*_1_, *d*_2_]. For a vertex *v* ∈*V*, let *a*(*v, l*) be a boolean value which is true if and only if there is a path of length *l* from source *s* to *v*. Then, the following recurrence solves the problem:

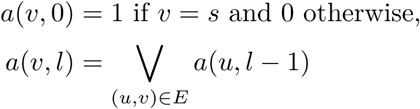

Solving the above recurrence requires filling a |*V*| × *d*_2_ table in column-wise order. The distance constraint from the source vertex *s* to *t* ∈ *V* is satisfied if and only if *a*(*t, l*) = 1 for any *l* ∈ [*d*_1_, *d*_2_]. Note that it is sufficient to store two columns in memory to fill the table, and an additional column to track the final result. The algorithm is summarized as a lemma below.

#### Lemma 3

*There exists an O*(*d*_2_|*E*|) *time and O*(|*V* |) *space algorithm that decides existence of a path of length d ∈* [*d*_1_, *d*_2_] *from one vertex to another in G*(*V, E*).

The time complexity of the above algorithm is significantly high, as it requires *O*(*d*_2_ |*E*|) time to validate distance constraints from a fixed source vertex. With some optimizations however, the above algorithm can be accelerated. As observed by Salmela *et al.* [39] in the context of gap-filling problem, we expect *d*_2_ ≪ |*V*|, therefore, it should be possible to compute a sub-graph containing vertices within ≤*d*_2_*/*2 distance from *v*_1_ or *v*_2_, before solving the recurrence. While this strategy was shown to be effective for gap-filling between assembled contigs, the count of vertex pairs to evaluate during read mapping process is expected to be significantly higher for large read sets. Reference genomes (e.g., GRCh38 for human genome) or graphs are static, or evolve slowly, in genomic analyses. As such, it is desirable to use an index-based strategy, where we pay a one-time cost to build an index, and validate the paired-end distance constraints quickly.

## 4 An Index-based Polynomial-time Algorithm

In the following, we describe our index construction and querying algorithm. Given a sequence graph in the form of a boolean adjacency matrix, the index construction procedure uses boolean matrix additions and multiplications. As we will note later, the worst-case time of building our index is polynomial in the input size, but still computationally prohibitive to handle real data instances. Subsequently, we will show how to exploit sparsity in graphs to accelerate the computation. The construction algorithm relies on the following boolean matrix operations.

### Definition 4

*Boolean matrix operations: Let A and B be two boolean n* ×*n matrices. The standard boolean matrix operations are evaluated in the following way:*

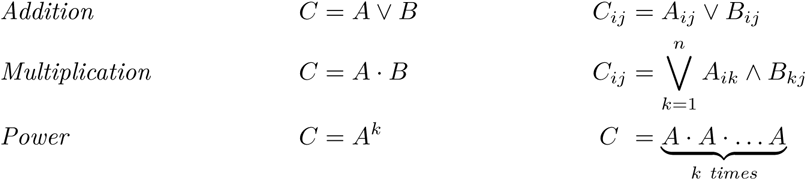

The boolean matrix addition and multiplication can also be performed using the standard matrix addition and multiplication, respectively. This is done by adjusting the non-zero values in output matrix to 1. Next, we define index matrix 𝒯, built using the adjacency matrix of the input graph and the distance parameters *d*_1_ and *d*_2_. Lemma 6 and 7 include its correctness proof and worst-case construction time complexity.

### Definition 5

*Let Adj be the* |*V* |×|*V* |*-sized boolean adjacency matrix associated with graph G*(*V, E*). *Define index matrix* 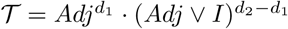, *where I is an identity matrix.*

### Lemma 6

𝒯 [*i, j*] = 1 *if and only if there exists a path of length d* ∈ [*d*_1_, *d*_2_] *from vertex v*_*i*_ *to vertex v*_*j*_.

**Proof.** Note that *Adj*^*k*^[*i, j*] = 1 if and only if there is a path of length *k* from vertex *v*_*i*_ to *v*_*j*_. To validate the paired-end distance constraints, we require 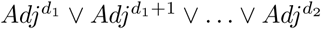.

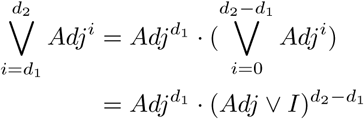

◂

### Lemma 7

*If multiplying two square matrices of dimension* |*V*| × |*V*| *requires O*(|*V*| ^*ω*^) *time, ω*∈ ℝ, *then computing the index matrix requires O*(|*V*| ^*ω*^ log(*d*_2_)) *time and O*(|*V*| ^2^) *space.*

**Proof.** Matrix addition uses *O*(|*V* |^2^) operations. Computing *A*^*k*^ requires *O*(|*V* |^*ω*^ log *k*) operations. Therefore, computing the index matrix requires *O*(|*V* |^2^ + |*V* |^*ω*^ (1 + log *d*_1_ + log(*d*_2_ *- d*_1_))) operations. As *ω ≥* 2 and *d*_2_ *≥ d*_1_, this simplifies to *O*(|*V* |^*ω*^ *·* log(*d*_2_)) time. ◂

The current best algorithm to compute general matrix multiplication requires *O*(|*V*| ^2.37^) time [23]. Using the general matrix storage format, querying for the distance constraints between a vertex pair requires a simple *O*(1) lookup. However, general matrix multiplication solvers require at least quadratic time and space (in terms of |*V*|), which does not scale to real graph instances. We next propose an alternate approach to build the index matrix that exploits sparsity in sequence graphs.

### 4.1 Exploiting Sparsity in Sequence Graphs

Typically, sequence graphs representing variation or assembly graphs have large diameter and high sparsity, with edge to vertex ratio close to 1 [31]. As *d*_2_ ≪ |*V*| in practice, we also expect our final index matrix to be sparse. As a result, we propose using SpGEMM (sparse matrix-matrix multiplication) operations to build the index. Below, we briefly recall the algorithm and matrix storage format used for SpGEMM. Subsequently, we shed light on the construction time and size of the index using this approach. We borrow standard notations typically used to discuss SpGEMM algorithms. Let *nnz*(*A*) denote number of non-zero values in matrix *A*. During boolean matrix multiplication *A · B*, let *bitops*(*AB*) indicate the count of non-zero bitwise-AND operations (i.e., 1 ∧ 1), assuming Definition 4.

#### 4.1.1 Working with Sparse Matrices

##### Storage

During SpGEMM, the input and output matrices are stored in a sparse format, such that the space is primarily used for non-zeros. Compressed Sparse Row (CSR) is a classic data structure for this purpose [3]. In CSR format, a boolean matrix *A*_*n*×*n*_ can be represented by using two arrays: the first array *ptr* of size *n* + 1 contains row pointers, and the second array *cols* of size *O*(*nnz*(*A*)) contains column indices of each non-zero entry in *A*, starting from the first row to the last. The row pointers are essentially offsets within the second array, such that the range *cols*[*ptr*[*i*]], *cols*[*ptr*[*i* + 1]] lists column indices in row *i*. By default, CSR format does not require the indices of a row to be sorted. However, the sorted order will be useful in our index storage to enable fast querying. Therefore, we use ‘sorted-CSR’ format in our application.

###### Remark 8

Storing a matrix *A*_*n*×*n*_ in sorted-CSR format requires Θ(*n* + *nnz*(*A*)) space.

###### Remark 9

Given a sequence graph *G*(*V, E*) as an array of edge tuples, transforming its adjacency information into sorted-CSR format takes Θ(|*V* | + |*E*|) time using count sort [15].

##### Multiplication (SpGEMM)

SpGEMM algorithms limit their operation count to just non-zero multiplications and additions required to compute the product, as the remaining entries are guaranteed to be 0. Most of the sequential and high-performance parallel algorithms for SpGEMM, including in MATLAB [13], are based on Gustavson’s algorithm [15]. The algorithm can take input matrices *A* and *B* in sorted-CSR format and produce the output matrix *C* = *A*·*B* in the same format. In this algorithm, a row of matrix *C*, i.e., *C*[*i*,:] is computed as a linear combination of the rows *κ* of *B* for which *A*[*i, κ*] ≠ 0. The complexity result from Gustavson’s work is listed as the following lemma.

###### Lemma 10

*The time complexity to multiply two sparse matrices A*_*n*×*n*_ *and B*_*n*×*n*_ *using Gustavson’s algorithm is* Θ(*n* + *nnz*(*A*) + *bitops*(*AB*)).

#### 4.1.2 Indexing Time and Storage Complexity

Computing the index (Definition 5) requires several SpGEMM operations. As such, it is hard to derive a tight bound on the complexity, as runtime and index size depend on non-zero structure of the input sequence graph. However, it is important to get an insight into how the different parameters, e.g., |*V*|, *d*_1_, *d*_2_ may affect them. To address this, we derive a practically useful lower-bound on the complexity.

Consider the chain graph *G*^′^(*V* ^′^, *E*^′^) associated with a longest path in *G*(*V, E*), *V* ^′^⊂ *V, E*^′^⊂ *E*. We claim that the time needed to index *G*(*V, E*) is either the same or worse than indexing its chain *G*^′^(*V* ^′^, *E*^′^) (Lemma 11). Subsequently, we compute the time complexity for indexing the chain using our SpGEMM-based algorithm. The rationale for analyzing the chain graph is (a) non-zero structure of a chain is simple and well-defined for computing time complexity, and (b) sequence graphs are expected to have ‘near-linear’ topology in practice, therefore the derived lower-bounds will be a useful indication of the true costs.

##### Lemma 11

*The time requirement for indexing a graph G*(*V, E*) *using SpGEMM is either the same or higher than indexing the chain graph G*^′^(*V* ^′^, *E*^′^) *associated with its longest path.*

**Proof.** Note that *G*^′^ is a sub-graph of *G*, which will also reflect in their adjacency matrices. The above lemma is based on the following simple observation. Suppose *A, B, A*^′^, *B*^′^, *δ*_1_, *δ*_2_ are boolean square matrices, such that, *A* = *A*^′^*V δ*_1_ and *B* = *B*^′^ *V δ*_2_. Then, using Gustavson’s algorithm, multiplying *A* and *B* requires at least as much time as required for multiplying *A*^′^ and *B*^′^. In addition, the product (*A · B*) is of the form (*A*^′^ *B*^′^) *V δ*_3_, where *δ*_3_ is a boolean matrix. For each SpGEMM executed while computing the index, we can use this argument to support the claim. ◂

##### Lemma 12

*Computing the index for G*(*V, E*) *using SpGEMM requires* Ω(|*V′*| ((*d*_2_ − *d*_1_)^2^ + log *d*_1_)) *time.*

**Proof.** Let *Adj*^*′*^ be the adjacency matrix associated with *G*^*′*^(*V* ^*′*^, *E*^*′*^). To prove the above claim, it is useful to visualize the structure of *Adj*^*′*^ (Figure 2). Define a constant *k* ≪|*V*| ^*′*^. For simplicity, assume *d*_1_ and *d*_2_ − *d*_1_ are powers of 2. Throughout the index computation, the time required by Gustavson’s SpGEMM algorithm is dictated by *bitops*. Following Lemma 10, multiplying *Adj*^*′k*^ to *Adj*^*′k*^ requires Θ(|*V* ^*′*^|) time. In addition, multiplying (*Adj*^*′*^ *V I*)^*k*^ with (*Adj*^*′*^ *V I*)^*k*^ requires Θ(|*V* ^*′*^|(*k*^2^)) time. Therefore, we need 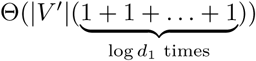 time to compute 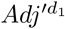, and Θ(|*V*′ | (1 + 2 ^2^+ 4^2^ + + …(*d*_2_ *- d*_1_)^2^)) time to compute 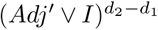. The final multiplication between 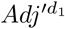 and 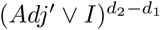 uses Θ(|*V* ^′^|(*d*_2_ *- d*_1_)) time. All these operations add up to Θ(|*V* ^′^|((*d*_2_ *- d*_1_)^2^ + log *d*_1_)) time. This argument, and Lemma 11 suffice to support the claim. ◂

**Figure 2.**
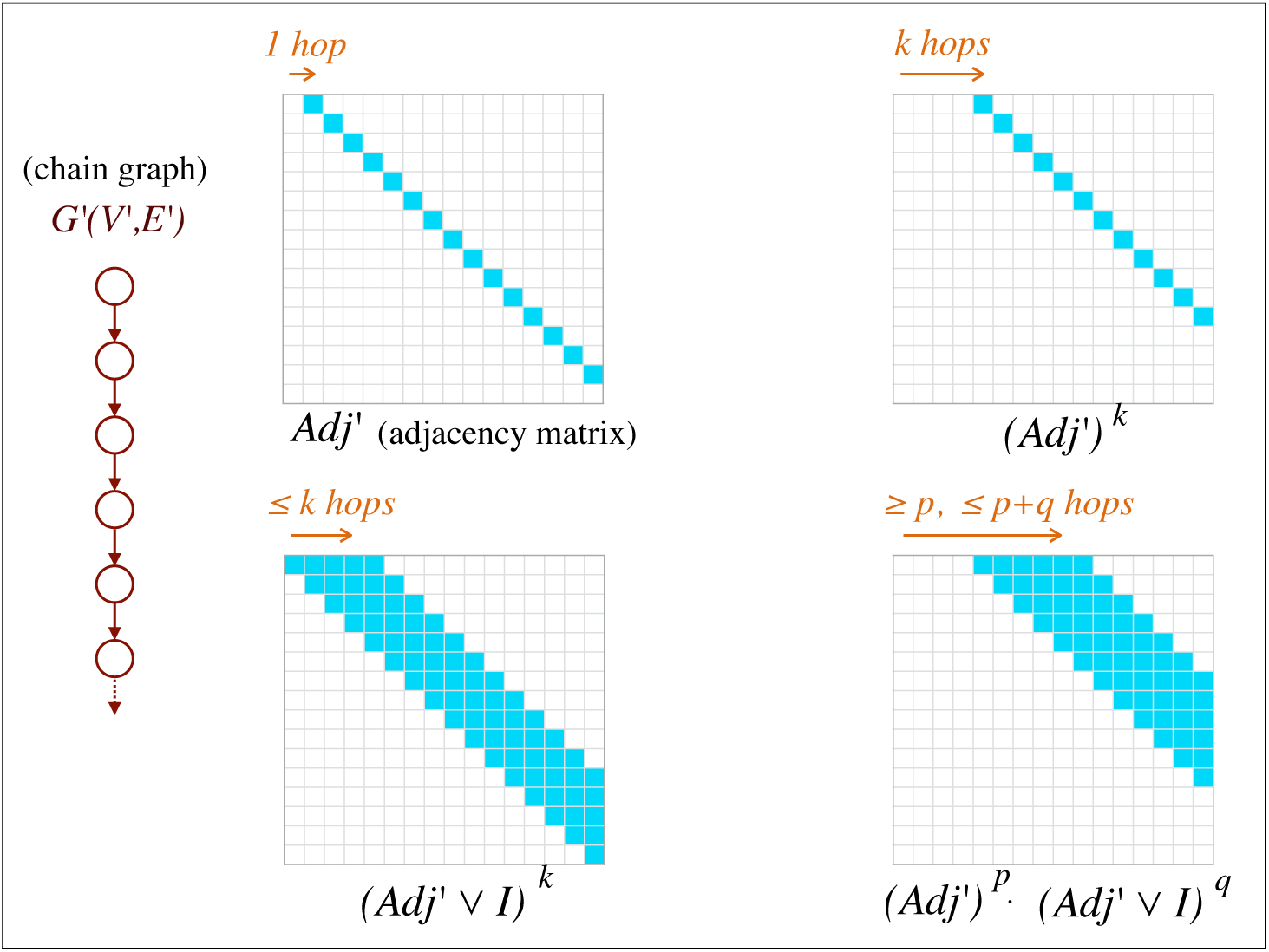
Visualizing non-zero structure of adjacency matrix of a chain graph. We also show how the structure changes after exponentiation. This is useful to count *bitops* during SpGEMM.

##### Remark 13

The index size is dictated by count of non-zeros in the final output matrix. Using similar arguments as above, it can be shown that the index storage for graph *G*(*V, E*) requires Ω(|*V* ^′^|(*d*_2_ *- d*_1_ + 1)) space.

#### 4.1.3 Querying the Index

Querying for a value in a sorted-CSR formatted index is trivial. The lookup procedure for *𝒯* [*i, j*] requires a binary search among the non-zeros of row *i*. Let *maxRownnz*(*𝒯*) be the maximum row size, i.e., maximum number of non-zero entries in a row of index matrix *𝒯*. After computing the index, deciding the existence of a path of length *d ∈* [*d*_1_, *d*_2_] between two vertices in *G*(*V, E*) requires *O*(log *maxRownnz*(*𝒯*)) time.

## 5 Results

We implemented our algorithm, referred to here as PairG, in C++. The source code is available at https://github.com/ParBLiSS/PairG. We conducted our evaluation using an Intel Xeon CPU E5-2680 v4, equipped with 28 physical cores and 256 GB main memory. In our implementation, we utilize KokkosKernels [9], an open-source parallel library for basic linear algebra (BLAS) routines. This library does not provide explicit support for boolean matrix operations, so we used integer matrix operations instead, while rounding the non-zero output values to one. We leveraged multi-threading support in KokkosKernels, and allowed it to use 28 threads during execution. Our benchmark data sets, summarized below, consist of cyclic and acyclic graphs built using publicly available real data. We tested indexing and querying performance using PairG for various values of distance constraints. These choices were motivated by the typical insert sizes used for Illumina paired-end sequencing. We demonstrate that PairG can index graphs with more than a million vertices in a reasonable time. Once the index is built, it can answer a million distance constraint queries in a fraction of a second.

### 5.1 Datasets

We generated seven sequence graphs, four acyclic (G1-G4) and three cyclic (G5-G7) (see Table 1). The first four sequence graphs are variation graphs built using human genome segments (GRCh37) and variant files from the 1000 Genomes Project (Phase 3) [6]. We used vg [12] for building these graphs. The human genomic regions considered in our evaluation are mitochondrial DNA (mtDNA), BRCA1 gene, the killer cell immunoglobulin-like receptors (LRC_KIR), and the major histocompatibility complex (MHC). The sizes of these regions range from 16.6 kilobases (mtDNA) to 5.0 megabases (MHC) in the human genome. De Bruijn Graph (DBG) is another popular format to represent ‘pan-genome’ of a species. DBG is also a good candidate structure to test our algorithm on more complex graphs. We built DBGs with *k*-mer length 25 using whole-genome sequences of one (G5), five (G6), and twenty (G7) B. *anthracis* strains, using SplitMEM [27]. Strain ids and size of these genomes are listed in the Appendix.

**Table 1.** Directed sequence graphs used for evaluation. In these graphs, each vertex is labeled with a DNA nucleotide. Four acyclic graphs are derived from segments of human genome and variant files from the 1000 Genomes Project (Phase 3). Three cyclic graphs are de Bruijn graphs built using whole-genome sequences of *Bacillus anthracis* strains, with *k*-mer length 25.

| Id | Graph | # vertices | # edges | Type |
| --- | --- | --- | --- | --- |
| G1 | mitochondrial-DNA | 21,038 | 26,842 | acyclic |
| G2 | BRCA1 | 83,268 | 85,422 |  |
| G3 | LRC_KIR | 1,108,769 | 1,154,046 |  |
| G4 | MHC | 5,138,913 | 5,318,639 |  |
| G5 | *B. anthracis* (1 strain) | 5,174,432 | 5,175,000 | cyclic |
| G6 | *B. anthracis* (5 strains) | 10,360,296 | 10,363,334 |  |
| G7 | *B. anthracis* (20 strains) | 11,237,067 | 11,253,437 |  |

We tested PairG using three different ranges of distance constraints, associated with three insert-size configurations. Assuming a typical sequencing scenario of insert-size configuration as 300 bp and read length 100 bp, the inner distance between paired-end reads should equal 100 bp (i.e., insert size minus twice the read length). To allow for sufficient variability, we tested PairG using *d*_1_ = 0, *d*_2_ = 250. Similarly, for insert-size configurations of 500 bp and 700 bp, we tested PairG using inner distance limits (*d*_1_ = 150, *d*_2_ = 450) and (*d*_1_ = 350, *d*_2_ = 650), respectively. There may be insert size configurations where allowing read overlaps may also be necessary; this can be handled trivially by computing a second matrix with desirable limit on the overlap length.

### 5.2 Index construction

We report wall-clock time, memory-usage, and index size (*nnz*) using various graphs and distance constraints in Table 2. We note large variation in performance for various graph sizes and complexity. Time, memory, and index size increase almost linearly with increasing graph size. This is also expected based on our theoretical analysis (Section 4.1.2). The specified range of distance constraints [*d*_1_, *d*_2_] can also affect performance. The index size for graphs G1-G5 remains almost uniform for the three different constraints. This should be because the first five graphs have linear chain-like topology, where the performance should be dictated by the gap (*d*_2_ − *d*_1_) according to our analysis. The graphs G1-G4 were built by augmenting the variations (substitutions, indels) on the reference sequence, and the fifth graph uses a single strain. On the other hand, index size varies with distance constraints for the last two graphs, especially G7. We expect G7 to have significantly more branching due to complex structural variations and repeats. Another important contributing factor is that DBGs collapse repetitive *k*-mers into a single vertex, causing large deviation from the linear topology. As a result, it takes time ranging from 1.3 hours to 17 hours for graph G7.

**Table 2.** Performance measured in terms of wall-clock time and memory-usage for building index matrix using all input graphs and different distance constraints. *nnz* represents number of non-zero elements in the index matrix, to indicate its size. Our implementation uses 4 bytes to store each non-zero of a matrix in memory.

| Id | Distance constraints |  |  |  |  |  |  |  |  |
| --- | --- | --- | --- | --- | --- | --- | --- | --- | --- |
|  | [0 - 250] |  |  | [150 - 450] |  |  | [350 - 650] |  |  |
|  | Time | Mem | *nnz* | Time | Mem | *nnz* | Time | Mem | *nnz* |
| G1 | 0.2s | 0.1G | 6.8M | 0.4s | 0.2G | 7.8M | 0.4s | 0.2G | 7.6M |
| G2 | 0.4s | 0.3G | 21M | 0.9s | 0.5G | 26M | 0.9s | 0.5G | 26M |
| G3 | 5.6s | 3.8G | 0.3B | 12s | 6.2G | 0.3B | 12s | 6.2G | 0.3B |
| G4 | 25s | 17G | 1.3B | 53s | 28G | 1.6B | 53s | 28G | 1.6B |
| G5 | 25s | 17G | 1.3B | 54s | 28G | 1.6B | 54s | 28G | 1.7B |
| G6 | 52s | 35G | 2.8B | 2m | 60G | 3.6B | 2m | 60G | 4.2B |
| G7 | 1.3h | 56G | 4.6B | 7.2h | 118G | 8.9B | 17.2h | 129G | 14B |

For larger graphs, we expect the index size (*nnz*) to increase with graph size. In sorted-CSR storage format, we use 4 bytes for each non-zero, which can become prohibitively large at the scale of complete human genome. However, we expect substantial room for compressing the final index, and plan to explore it in the future. Memory-usage of a matrix operation (addition or multiplication) is dictated by the size of the associated input and output matrices. As a result, memory required for the index construction appears to increase proportionally with the index size (Table 2).

### 5.3 Querying Performance

While indexing is a one-time routine for a sequence graph, we would need to query the index numerous times during the mapping process. Querying for a vertex pair requires a simple and fast lookup in the index (Section 4.1.3). As read mapping locations are expected to be uniformly distributed over the graph, we tested the querying performance by generating a million random vertex pairs (*u, v*), *u, v* ∈ [1, | *V* |]. For all the seven graphs, querying a million vertex pairs finished in less than a second (Table 3). Even though the majority of randomly generated queries result in a ‘no’ answer, this aspect has insignificant effect on the query performance. Our results implicate that distance constraints can be validated exactly without additional overhead on the mapping time, which is similar to the case of mapping reads to a reference sequence.

**Table 3.** Time to execute a million queries using all the graphs and distance constraints. Each query is a random pair of vertices in the graph.

| Id | Distance constraints |  |  |
| --- | --- | --- | --- |
|  | [0 - 250] | [150 - 450] | [350 - 650] |
|  | Time (sec) |  |  |
| G1 | 0.1 | 0.1 | 0.1 |
| G2 | 0.2 | 0.2 | 0.2 |
| G3 | 0.4 | 0.4 | 0.5 |
| G4 | 0.5 | 0.5 | 0.5 |
| G5 | 0.4 | 0.5 | 0.5 |
| G6 | 0.4 | 0.5 | 0.5 |
| G7 | 0.5 | 0.5 | 0.6 |

We also compared our index-based algorithm using a breadth first search (BFS)-based heuristic. Using this heuristic, we answer a query of a pair of vertices (*u, v*) as true if and only if *v* is reachable within *d*_2_ distance from *u*. Accordingly, BFS initiated from vertex *u* is terminated if we find vertex *v*, or after we have explored all vertices up to depth *d*_2_. This heuristic does not require a pre-computed index. However, we find that querying time using our index-based algorithm is faster by two to three orders of magnitude. For the randomly generated query set, fraction of results that agree between the two approaches varied from 98% to 100%. The above heuristic yields incorrect result for a vertex pair (*u, v*) when all the possible paths that connect *u* to *v* have length < *d*_1_.

## 6 Conclusions and Future Work

In this work, we formulated the Paired-end Validation Problem, required for paired-end read mapping on sequence graphs. We proposed the first provably good and practically useful exact algorithm for solving this problem. The proposed algorithm builds on top of existing SpGEMM algorithms, to exploit the sparsity and large diameter characteristics of sequence graphs. Our experiments indicate that index construction time is affected by the size and topology of the sequence graph, as well as the desired distance constraints. The querying time is less than a second for answering a million distance queries using all test cases.

There are two immediate avenues for future work. First, we expect huge room for lossless compression in the final index matrix. By computing an appropriate vertex ordering, we can potentially benefit from differential coding of column indices. Thus, alternative formats than sorted-CSR might be more efficient for index storage. We anticipate that good compression ratio can be achieved without affecting querying performance. Second, we plan to integrate this work with an exact sequence to graph aligner as a paired-end read mapping tool, and evaluate the advantages of a provably-good approach relative to existing heuristics-based methods.

## Acknowledgements

The authors thank Abdurrahman Yasar, Siva Rajamanickam and Srinivas Eswar for sharing their insights on sparse matrix manipulations.

## A Appendix: Data Availability

**Table 4.** List of 20 *Bacillus anthracis* strains used to build the sequence graphs G5-G7. We used the first strain in G5, the first five strains in G6, and all the 20 strains in G7.

| Strain id | Assembly id | Size (Mbp) |
| --- | --- | --- |
| Ames | GCA_000007845.1 | 5.23 |
| delta Sterne | GCA_000742695.1 | 5.23 |
| Cvac02 | GCA_000747335.1 | 5.23 |
| Parent1 | GCA_001683095.1 | 5.23 |
| PR09-1 | GCA_001683255.1 | 5.23 |
| Sterne | GCA_000008165.1 | 5.23 |
| CNEVA-9066 | GCA_000167235.1 | 5.49 |
| A1055 | GCA_000167255.1 | 5.37 |
| A0174 | GCA_000182055.1 | 5.29 |
| Sen2Col2 | GCA_000359425.1 | 5.17 |
| SVA11 | GCA_000583105.1 | 5.49 |
| 95014 | GCA_000585275.1 | 5.34 |
| Smith 1013 | GCA_000742315.1 | 5.29 |
| 2000031021 | GCA_000742655.1 | 5.33 |
| PAK-1 | GCA_000832425.1 | 5.40 |
| Pasteur | GCA_000832585.1 | 5.23 |
| PR06 | GCA_001683195.1 | 5.23 |
| 55-VNIIVViM | GCA_001835485.1 | 5.35 |
| Sterne 34F2 | GCA_002005265.1 | 5.39 |
| UT308 | GCA_003345105.1 | 5.37 |

## References

1 Stefano Beretta, Paola Bonizzoni, Luca Denti, Marco Previtali, and Raffaella Rizzi. Mapping RNA-seq data to a transcript graph via approximate pattern matching to a hypertext. In International Conference on Algorithms for Computational Biology, pages 49–61. Springer, 2017.

2 Alexander Bowe, Taku Onodera, Kunihiko Sadakane, and Tetsuo Shibuya. Succinct de bruijn graphs. In International Workshop on Algorithms in Bioinformatics, pages 225–235. Springer, 2012.

3 Aydin Buluç, John Gilbert, and Viral B Shah. Implementing sparse matrices for graph algorithms. In Graph Algorithms in the Language of Linear Algebra, pages 287–313. SIAM, 2011.

4 Stefan Canzar and Steven L Salzberg. Short read mapping: An algorithmic tour. Proceedings of the IEEE, 105(3):436–458, 2015.

5 Computational Pan-Genomics Consortium. Computational pan-genomics: status, promises and challenges. Briefings in bioinformatics, 19(1):118–135, 2016.

6 1000 Genomes Project Consortium et al. A global reference for human genetic variation. Nature, 526(7571):68, 2015.

7 Thomas H Cormen, Charles E Leiserson, Ronald L Rivest, and Clifford Stein. Introduction to algorithms. MIT press, 2009.

8 Luca Denti, Raffaella Rizzi, Stefano Beretta, Gianluca Della Vedova, Marco Previtali, and Paola Bonizzoni. Asgal: aligning RNA-Seq data to a splicing graph to detect novel alternative splicing events. BMC bioinformatics, 19(1):444, 2018.

9 Mehmet Deveci, Christian Trott, and Sivasankaran Rajamanickam. Performance-portable sparse matrix-matrix multiplication for many-core architectures. In 2017 IEEE International Parallel and Distributed Processing Symposium Workshops (IPDPSW), pages 693–702. IEEE, 2017.

10 Alexander Dilthey, Charles Cox, Zamin Iqbal, Matthew R Nelson, and Gil McVean. Improved genome inference in the MHC using a population reference graph. Nature genetics, 47(6):682, 2015.

11 Alexander Dilthey, Pierre-Antoine Gourraud, Alexander J Mentzer, Nezih Cereb, Zamin Iqbal, and Gil McVean. High-accuracy HLA type inference from whole-genome sequencing data using population reference graphs. PLoS computational biology, 12(10):e1005151, 2016.

12 Erik Garrison, Jouni Sirén, Adam M Novak, Glenn Hickey, Jordan M Eizenga, Eric T Dawson, William Jones, Shilpa Garg, Charles Markello, Michael F Lin, et al. Variation graph toolkit improves read mapping by representing genetic variation in the reference. Nature biotechnology, 2018.

13 John R Gilbert, Cleve Moler, and Robert Schreiber. Sparse matrices in MATLAB: Design and implementation. SIAM Journal on Matrix Analysis and Applications, 13(1):333–356, 1992.

14 Richard E Green, Johannes Krause, Adrian W Briggs, Tomislav Maricic, Udo Stenzel, Martin Kircher, Nick Patterson, Heng Li, Weiwei Zhai, Markus Hsi-Yang Fritz, et al. A draft sequence of the neandertal genome. science, 328(5979):710–722, 2010.

15 Fred G Gustavson. Two fast algorithms for sparse matrices: Multiplication and permuted transposition. ACM Transactions on Mathematical Software (TOMS), 4(3):250–269, 1978.

16 Mahdi Heydari, Giles Miclotte, Yves Van de Peer, and Jan Fostier. Browniealigner: accurate alignment of illumina sequencing data to de bruijn graphs. BMC bioinformatics, 19(1):311, 2018.

17 Zamin Iqbal, Mario Caccamo, Isaac Turner, Paul Flicek, and Gil McVean. De novo assembly and genotyping of variants using colored de bruijn graphs. Nature genetics, 44(2):226, 2012.

18 Chirag Jain, Sanchit Misra, Haowen Zhang, Alexander Dilthey, and Srinivas Aluru. Accelerating sequence alignment to graphs. In 2019 IEEE International Parallel and Distributed Processing Symposium (IPDPS). IEEE, 2019 (to appear).

19 Chirag Jain, Haowen Zhang, Yu Gao, and Srinivas Aluru. On the complexity of sequence to graph alignment. In Research in Computational Molecular Biology, pages 85–100, Cham, 2019. Springer International Publishing.

20 Vaddadi Naga Sai Kavya, Kshitij Tayal, Rajgopal Srinivasan, and Naveen Sivadasan. Sequence alignment on directed graphs. Journal of Computational Biology, 26(1):53–67, 2019.

21 Daehwan Kim, Joseph M Paggi, and Steven Salzberg. Hisat-genotype: Next generation genomic analysis platform on a personal computer. BioRxiv, page 266197, 2018.

22 Ben Langmead and Steven L Salzberg. Fast gapped-read alignment with bowtie 2. Nature methods, 9(4):357, 2012.

23 François Le Gall. Powers of tensors and fast matrix multiplication. In Proceedings of the 39th international symposium on symbolic and algebraic computation, pages 296–303. ACM, 2014.

24 Heng Li. Aligning sequence reads, clone sequences and assembly contigs with bwa-mem. arXiv preprint arXiv:1303.3997, 2013.

25 Antoine Limasset, Bastien Cazaux, Eric Rivals, and Pierre Peterlongo. Read mapping on de bruijn graphs. BMC bioinformatics, 17(1):237, 2016.

26 Bo Liu, Hongzhe Guo, Michael Brudno, and Yadong Wang. debga: read alignment with de bruijn graph-based seed and extension. Bioinformatics, 32(21):3224–3232, 2016.

27 Shoshana Marcus, Hayan Lee, and Michael C Schatz. Splitmem: a graphical algorithm for pan-genome analysis with suffix skips. Bioinformatics, 30(24):3476–3483, 2014.

28 Tom O Mokveld, Jasper Linthorst, Zaid Al-Ars, and Marcel Reinders. Chop: Haplotype-aware path indexing in population graphs. bioRxiv, 2018.

29 Martin D Muggli, Alexander Bowe, Noelle R Noyes, Paul S Morley, Keith E Belk, Robert Raymond, Travis Gagie, Simon J Puglisi, and Christina Boucher. Succinct colored de bruijn graphs. Bioinformatics, 33(20):3181–3187, 2017.

30 Gonzalo Navarro. Improved approximate pattern matching on hypertext. Theoretical Computer Science, 237(1-2):455–463, 2000.

31 Adam M Novak, Glenn Hickey, Erik Garrison, Sean Blum, Abram Connelly, Alexander Dilthey, Jordan Eizenga, MA Saleh Elmohamed, Sally Guthrie, André Kahles, et al. Genome graphs. bioRxiv, page 101378, 2017.

32 Esko Nuutila. Efficient transitive closure computation in large digraphs. Finnish Academy of Technology, 1998.

33 Matti Nykänen and Esko Ukkonen. The exact path length problem. Journal of Algorithms, 42(1):41–53, 2002.

34 Benedict Paten, Adam M Novak, Jordan M Eizenga, and Erik Garrison. Genome graphs and the evolution of genome inference. Genome research, 27(5):665–676, 2017.

35 Goran Rakocevic, Vladimir Semenyuk, Wan-Ping Lee, James Spencer, John Browning, Ivan J Johnson, Vladan Arsenijevic, Jelena Nadj, Kaushik Ghose, Maria C Suciu, et al. Fast and accurate genomic analyses using genome graphs. Technical report, Nature Publishing Group, 2019.

36 Mikko Rautiainen and Tobias Marschall. Aligning sequences to general graphs in O(V + mE) time. bioRxiv, 2017. URL: https://www.biorxiv.org/content/early/2017/11/08/216127.

37 Mikko Rautiainen, Veli Mäkinen, and Tobias Marschall. Bit-parallel sequence-to-graph alignment. Bioinformatics, 03 2019. doi:10.1093/bioinformatics/btz162.

38 David Reich, Michael A Nalls, WH Linda Kao, Ermeg L Akylbekova, Arti Tandon, Nick Patterson, James Mullikin, Wen-Chi Hsueh, Ching-Yu Cheng, Josef Coresh, et al. Reduced neutrophil count in people of african descent is due to a regulatory variant in the duffy antigen receptor for chemokines gene. PLoS genetics, 5(1):e1000360, 2009.

39 Leena Salmela, Kristoffer Sahlin, Veli Mäkinen, and Alexandru I Tomescu. Gap filling as exact path length problem. Journal of Computational Biology, 23(5):347–361, 2016.

40 Jouni Sirén. Indexing variation graphs. In 2017 Proceedings of the ninteenth workshop on algorithm engineering and experiments (ALENEX), pages 13–27. SIAM, 2017.

41 Jouni Sirén, Niko Välimäki, and Veli Mäkinen. Indexing graphs for path queries with applications in genome research. IEEE/ACM Transactions on Computational Biology and Bioinformatics (TCBB), 11(2):375–388, 2014.

